# Both terminal misfolding and polymerisation contribute to disease-relevant responses in cell models of α_1_-antitrypsin deficiency-associated liver disease

**DOI:** 10.1101/2025.09.30.679226

**Authors:** Panayiota Stylianou, Kish Adoni, Mun Peak Nyon, Adam Cryar, Imran Haq, Jennifer A. Dickens, Charis-Patricia Segeritz, Graeme C. Smith, Adriana Ordonez, James A. Irving, David A. Lomas, Kyra-Mae Leighton, Jessica Beasley, Aleck W.E. Jones, S. Tamir Rashid, Ludovic Vallier, Konstantinos Thalassinos, Bibek Gooptu

## Abstract

Polymerisation of α_1_-antitrypsin within hepatocytes is considered central to the pathogenesis of α_1_-antitrypsin deficiency-associated liver fibrosis, most commonly in homozygotes for the Z (p.Glu342Lys) allele. Polymerisation proceeds via self-association of monomeric intermediate states. In parallel, >50% of synthesised Z α_1_-antitrypsin is instead recognized as terminally-misfolded and degraded. It is unclear whether this contributes to Z α_1_-antitrypsin deficiency-associated liver disease. We characterised the relationships between polymer formation, terminal misfolding and their cellular consequences, using label-free proteomics mass spectrometry (MS), light and electron microscopy, and cellular assays. Proteomic analyses of well-established CHO cell models of hepatocyte handling of α_1_-antitrypsin variants indicated that cellular responses to the Z mutation were surprisingly similar to those seen with the Null_HongKong_ variant (NHK), which can only misfold terminally and cannot polymerise. A minor set of proteins showed increases associated with Z and not NHK α_1_-antitrypsin expression, consistent with a polymer-specific response, characterized by association with increased organellar organization and vesicle-mediated transport. Conversely, proteostatic and pro-fibrotic integrin-associated pathways increased with the degree of terminal misfolding of the expressed α_1_-antitrypsin variant. Bioenergetic pathway changes indicated concomitant switching from oxidative to glycolytic metabolism. Cell studies further correlated fibrosis-associated behaviours with terminal misfolding rather than polymerisation. Terminal misfolding, as well as polymerisation behaviour, may therefore be important for pro-fibrotic responses including metabolic reprogramming and senescence in Z α_1_-antitrypsin deficiency. Molecular therapies may prove most efficacious for associated liver disease if they address terminal misfolding as well as polymerisation.

## Introduction

The Z (Glu342Lys) variant of α_1_-antitrypsin confers a high risk of liver fibrosis (1) leading to prevalence of clinically-significant liver disease (e.g. cirrhosis, hepatocellular carcinoma) in 10% of homozygotes (2). The heterozygote state (PiMZ) is also associated with ∼8-fold increased risk of liver disease relative to wild-type (M) homozygotes (PiMM) (3). The risk arises due to toxic conformational behaviour of the mutant α_1_-antitrypsin polypeptide chain after translation, during and/or after folding within the hepatocyte endoplasmic reticulum (ER) (4–7). Only about 15% of synthesized Z α_1_-antitrypsin is secreted and this has impaired antiprotease function (8). The loss-of-function consequences within the lung, the organ where α_1_-antitrypsin function is critical, predispose to early-onset emphysema (9).

Hepatocytes from PiZZ individuals with liver disease are characterized by ER inclusion bodies composed of α_1_-antitrypsin molecules, non-covalently linked into polymeric chains that entangle and precipitate (7). Other α_1_-antitrypsin variants associated with deficiency are also polymerogenic (10–14). Where assessed, polymerisation propensity generally correlates with severity of disease phenotype, and inversely with ability to fold to the native state associated with inhibitory function. However, in the case of the Baghdad variant, refolding studies indicated it both polymerizes more readily yet folds better than Z α_1_-antitrypsin (15). Polymerisation and terminal misfolding, while related processes, may therefore give rise to distinct pools of conformational behaviour (Supp. Fig. 1).

**Fig. 1:**
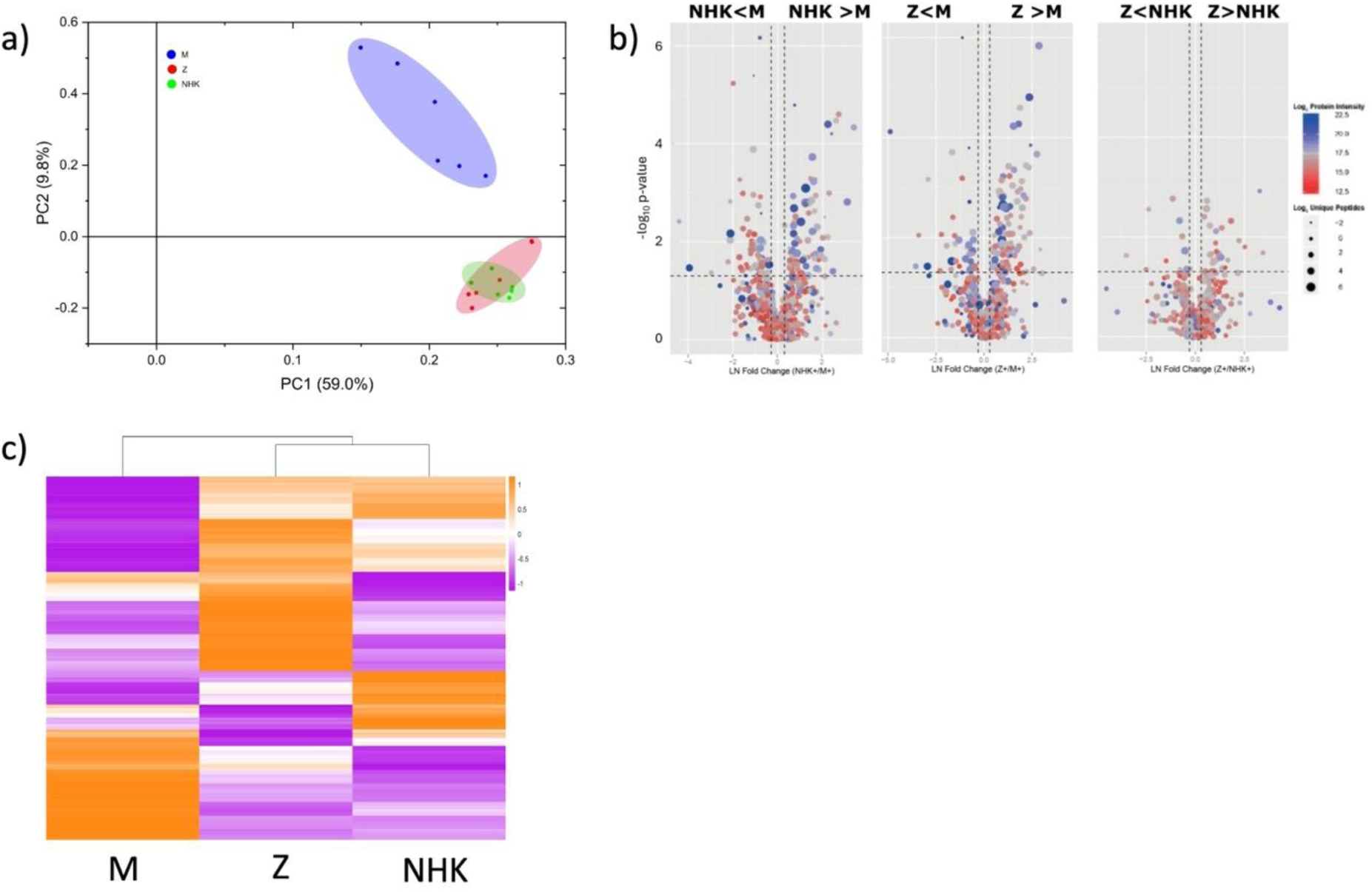
Proteomic analysis of CHO cells expressing M, Z or NHK α_1_-antitrypsin demonstrates high global similarity for expression of Z and NHK variants relative to. **M** a) Principal component (PC) analysis. b) Volcano plot, vertical dashed lines indicate threshold for confident discrimination of change (log_2_-fold change=1.3), and horizontal dashes delineate the p=0.05 probability threshold following Benjamini-Hochberg correction for multiple comparisons. Each point plotted represents an identified protein, with log_2_ mapping of the number of unique peptides used for its assignment (spot radius), and of signal intensity (red-to-blue heatmap colouring as indicated). c) Hierarchical cluster analyses (low-to-high detected levels are depicted in purple-to-orange spectrum as indicated).

In PiZZ individuals, polymer levels within liver cells and extracellularly within the circulation, appear to correlate with key disease features (16, 17). Moreover, polymerogenic mutants of related proteins within the serine protease inhibitor (serpin) superfamily cause a range of diseases (serpinopathies) through analogous combinations of gain- and loss-of-function effects (18). ∼15% of synthesized α_1_-antitrypsin is retained within the polymeric inclusions and >50% is recognized as terminally misfolded and targeted for ER associated degradation (ERAD) (4, 6). Polymer accumulation, however, does not trigger detectable unfolded protein (UPR) and ER stress responses *per se* in cell models of disease-mutant α_1_-antitrypsin expression that recapitulate *in vivo* hepatocyte phenotypes, though it is associated with sensitisation to this (19). A separate ‘ordered protein response’ has been observed in cell models of α_1_-antitrypsin deficiency (20). Polymers are also secreted extracellularly and have pro-inflammatory effects (21–23). Together these findings have led to a consensus model of α_1_-antitrypsin deficiency pathogenesis where polymerisation is regarded as a central event (24). On this basis we and others have worked to develop small molecules specifically designed to block the conformational changes that lead to polymerisation (25–31). Null variants of α_1_-antitrypsin, where α_1_-antitrypsin is not detectable by standard biochemical assays within the hepatocyte or in the circulation, are believed to contribute to substantial risk of emphysema but not to liver disease (32).

There is a need to develop a better understanding of the mechanistic steps by which the aberrant conformational events associated with folding of the Z and other common, polymerogenic variants of α_1_-antitrypsin within the ER trigger hepatic fibrosis. We previously used transcriptomic and proteomic techniques to study the effects of the Z mutation of α_1_-antitrypsin in the inducible pluripotent stem cell (iPSC)-derived hepatocyte-like (iHep) cell model of disease derived from a patient and validated our findings in PiZZ liver disease (33). However, this was not cross-referenced with a model of terminal misfolding alone. Changes associated with polymerisation *per se* could therefore not be distinguished from those related to terminal misfolding and high activity of the ERAD system.

To address this, we performed proteomic studies in the well-characterised and validated CHO cell model of hepatocyte behaviour in α_1_-antitrypsin deficiency. This allowed us to compare findings in the contexts of stable transfection and inducible expression of M and Z α_1_-antitrypsin but also with expression of the rarer Null_HongKong_ (NHK) variant where a point mutation at residue 318 introduces 15 new residues and a 61 amino-acid C-terminal truncation (34). These changes result in terminal misfolding of 100% of synthesized α_1_-antitrypsin polypeptide chains, saturating the ERAD response and triggering the UPR and ER stress. Indeed, NHK α_1_-antitrypsin expression is widely used as an experimental trigger in studies characterising these responses (34–39). Expression of the Z variant of α_1_-antitrypsin does not cause ER stress in the CHO cell model and similar findings have been reported in human hepatocytes. However, increased susceptibility to ER stress is observed in these cells in the context of further experimental challenge. We have proposed a “two hit” model for ER stress in response to Z α_1_-antitrypsin expression, and that this susceptibility relates to the terminally-misfolded polypeptide fraction reducing reserve proteostatic capacity within the ER. We hypothesized that the toxic gain-of-function pathway attributable to Z α_1_-antitrypsin polymer fraction would be defined by a proteomic signature distinct from those seen with either expression of the wild-type (M) or the NHK variant of α_1_-antitrypsin.

## Materials and Methods

### Preparation and proteomics of CHO cell ER samples

Chinese hamster ovary (CHO)-K1 cells stably transfected with Tet-On expression systems for M, Z, or Null_HongKong_ (NHK) variants of α_1_-antitrypsin were cultured, and expression induced with doxycycline, as described previously (19). Cells were harvested and lysed after 5 days post-seeding (2 days post-induction). Lysates were fractionated and processed for GeLC-MS^E^ as described previously (40) LC-MS^E^ data were processed using ProteinLynx Global Server (PLGS) v2.5.2. The data processing prior to database searching included ion detection, alignment and clustering methods detailed elsewhere (41). Thresholds used for peak selection were as follows, 150 counts for low energy ion detection, 40 counts for high energy ion detection and 500 counts for precursor exact mass retention time (EMRT) integrated intensity. Samples were run on a Waters Synapt G2 with a Waters Acquity LC system.

### Database searches

Data were searched using PLGS v2.5.2 against a Chinese Hamster Ovary (CHO) database (www.chogenome.org/files/CHO_refseq_protein.fasta) appended with human α_1_-antitrypsin and common contaminant proteins. Carbamidomethyl (C) was set as a fixed modification while oxidation (M), acetylation (K & N-term) and deamidation (N &Q) were set as variable modifications. Precursor and fragment ion tolerances were determined automatically by PLGS. Protein identification criteria were set as the detection of at least 2 fragments ion per peptide, at least 2 fragments per protein and at least 1 peptide identification per protein. A maximum of 2 missed cleavages were allowed. The protein level false discovery rate was maintained at < 5%, estimated based upon the number of proteins identified from a randomized database. Post-processing the false discovery rate was minimized further by applying the additional criterion that for a successful identification a protein must be identified in a minimum of two biological repeats. The mass spectrometry data and associated database search results have been deposited to the ProteomeXchange Consortium via the PRIDE partner repository with the dataset identifier PXD002219 and 10.6019/PXD002219.

### Protein quantification

Protein quantification was achieved through the Hi3 technique. Hi3 is based upon the observation that the summed intensity of the three most ionizing peptides per protein correlates with high accuracy and precision to protein concentration (42). As label-free quantification directly compares samples analysed in concurrent runs, quantification errors can be introduced by run-to-run instrument variability. To reduce this; within-condition normalization was applied prior to quantification. This was undertaken at the peptide level using a script written in-house in R to implement locally-weighted scatter point smoothing (LOESS, Fig. 1d). Post LOESS normalization, further global normalization was applied based upon the total detected protein intensity per run, in order to account for sample preparation and loading variability across biological replicates and across conditions.

### Differential expression

Pairwise comparisons of relative peptide abundances were performed for the three α_1_-antitrypsin expression contexts. Differential abundance was calculated based on fold change and statistical analysis. Fold change was calculated for each comparison using the mean of Hi3 protein intensity across replicates. P values were calculated using a two-sided Student’s t-test, with equal variance and performed on log_2_ transformed intensity data. Benjamini-Hochberg correction was applied for multiple comparisons. Proteins that were calculated to have a log_2_ fold change larger than 1.3 and a p value < 0.05 were defined as differentially expressed. MS^E^ quantitative technical variation has been previously demonstrated to be reproducible at 10-15% (40). A 1.3x log_2_ fold threshold therefore represents a value typically 2-3 times larger than estimated technical measurement error and compares well with more established label-based methods.

### Data Processing

For quantitative analysis of proteomics data across the three conditions including control (M), Z and NHK variant CHO cells; data were filtered such that each protein must be identified in at least 1 biological replicate (and two technical replicates) of each of the three conditions. As such, proteins that were not detected in all three conditions were excluded from further analysis. IBAQ values were imputed for missing values as an average of the identified IBAQ values of that condition. Principle component analysis (PCA) was performed using PC1 (59.0%) and PC2 (9.8%) with Origin v2024b (OriginLab Corporation, Northampton, MA, USA) using the parameter settings analyze: Correlation Matrix, Number of Components to Extract: 2. Hierarchical Clustering Analysis was also performed using Origin v2024b with the mean IBAQ value of the protein for each condition (for columns and rows; Cluster Method: Group Average, Distance Type: Pearson Correlation, Number of Clusters: 1). Next, proteins were grouped according to the following IBAQ value criteria: NHK>Z>M, M>Z>NHK, Z<M<NHK and Z<NHK<M. For each criterion, the protein list was analysed using STRING v12.0 (43) with standard settings (network edges: confidence, all active interaction scores, confidence: 0.4). Networks were then exported to Cytoscape v3.10.3 (44) where functional analysis was performed for biological process ontology and network annotations. For investigation of specific pathways, all validated proteins were input into STRING and the list of proteins associated with specific Gene Ontology Biological Processes were extracted. The log_2_ IBAQ of the proteins, in conditions M, Z and NHK were then visualised via a heatmap using Prism GraphPad v10.4.2.

### Cell phenotyping

To assess mitochondrial morphology, cultured CHO-K1 cells expressing M, Z or NHK variants of α_1_-antitrypsin mutation were incubated with the MitoTracker Red stain M7512 Red CMXRos (Life Technologies, ThermoFisher, MA, USA) as per manufacturer’s instructions. Live cells were imaged on an incubated LSM710 laser point scanning confocal microscope (Carl Zeiss Ltd, Cambridge UK) and acquired with the Zen 2010 software using a 63x 1.4NA Plan Apochromat objective lens. The experiments were repeated 3 times for each cell line. For quantitative analysis of mitochondrial morphology, mitochondria were selected in an automated manner and their sphericity scores calculated (Volocity, PerkinElmer, MA, USA; Imaris, Bitplane, Zurich, Switzerland).

Senescence behaviour was evaluated by assays to quantify senescence associated-β-galactosidase expression, proliferation by BrdU uptake, and p16 expression as described previously (45, 46). Immunofluorescence studies were undertaken using antibodies against KDEL motif (ENZ-ABS679, Enzo Life Sciences), Drp1 and Mfn-1 (respectively D6C7 #8570 and D6E2S #14739, Cell Signaling Technology), galectin-3 (sc-23938 produced by Santa Cruz Biotechnology, TX, USA), β1-integrin (ab24693, Abcam, Cambridge, UK) and p16 (ab241543, Abcam, Cambridge UK). Cells were immunostained (46) and imaging was undertaken on a Leica TCS SP5 microscope.

## Results

### Global proteomic profiles indicate high similarity in cell responses to Z and NHK variant relative to M α_1_-antitrypsin expression

We explored the proteomic profiles of existing cell models for the expression of M, Z and NHK α_1_-antitrypsin that recapitulate their expression and subcellular behaviour in hepatocytes. Contrary to our initial hypothesis, both Z and NHK α_1_-antitrypsin expression were associated with similar global proteomic profiles relative to the M protein whether assessed by principal component analysis (Fig. 1a), volcano plot (Fig. 1b), or hierarchical cluster analysis (Fig. 1c). The data were interrogated to better understand the relationships between the observations, polymerisation and terminal misfolding.

### Cellular pathways most associated with polymerisation

To first explore the set of proteins whose relative abundance was most up- or down-regulated with polymerisation, we identified proteins whose abundance was greatest in cells expressing Z α_1_-antitrypsin and least in those expressing the NHK variant, and vice versa. These were termed NHK<M<Z and NHK>M>Z respectively (Fig. 2a). This strategy should down-prioritise responses associated with terminal misfolding. Relatively small subsets of proteins were identified in each case. Interestingly, the profile associated with the order NHK<M<Z was enriched for organelle organisation and vesicle mediated transport. This may reflect changes to ER organisation described in association with polymer accumulation. The profile defined NHK>M>Z represented very few proteins.

**Fig. 2:**
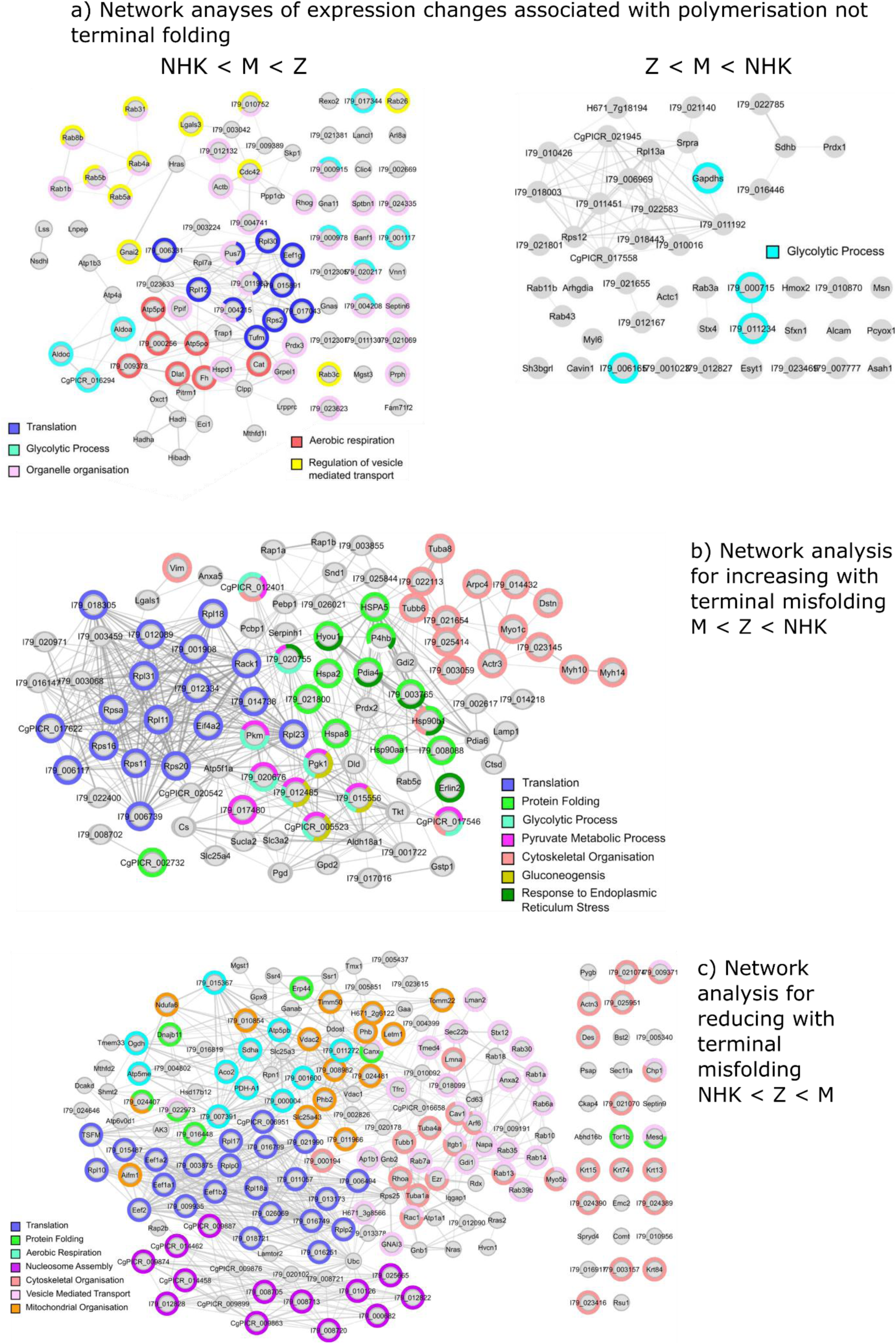
Network analyses of functional clusters identify distinct response repertoires to polymerisation and misfolding. Medium stringency STRING analyses of proteins differentially co-expressed with Z and NHK variants, relative to observations with expression of M α_1_-antitrypsin in CHO cells. a) Changes observed with increasing (left, NHK<M<Z) and decreasing (right, Z<M<NHK) ratios of polymerisation:ERAD behaviours of α_1_-antitrypsin variants. b) Protein increases observed with increasing terminal misfolding behaviours (M<Z<NHK). c) Protein increases observed with decreasing terminal misfolding behaviours (NHK<Z<M).

Proteins whose expression levels followed the order M<NHK<Z, or M>NHK>Z are shown in Supp. Fig. 2 and represent changes that are qualitatively similar for both variants relative to wild-type expression. Pathway analyses for all identified protein profiles are shown in Supp. Fig. 3-8.

**Fig. 3:**
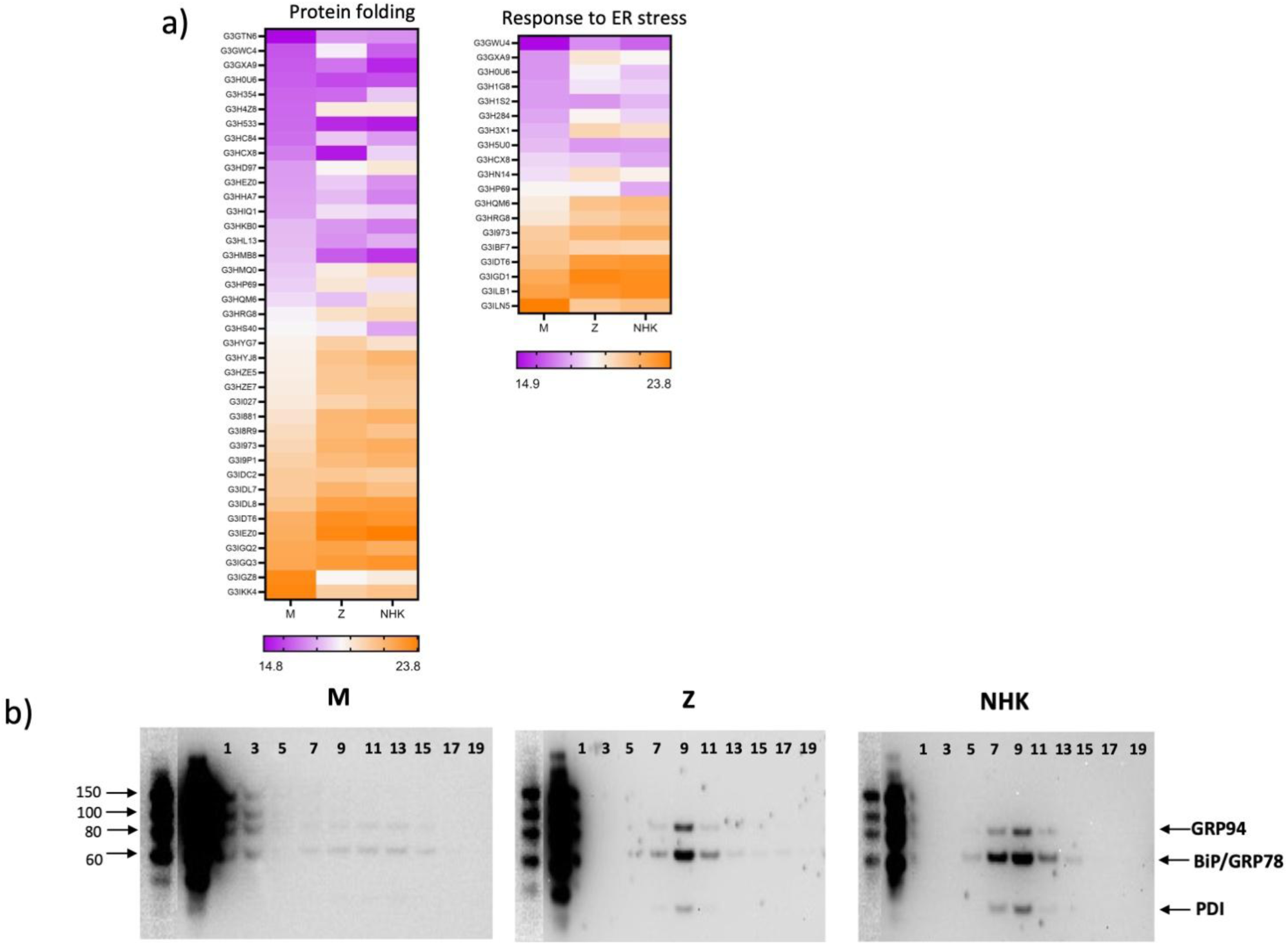
Increases in proteostatic response M<Z<NHK. a) Heatmap depiction of proteins associated with protein folding processes and ER stress response responses. Proteins are listed in order of increasing levels in cells expressing M α_1_-antitrypsin. Purple depicts the lowest levels detected and orange the highest. Thus in this case, both sets of proteins show generally increased co-expression with the α_1_-antitrypsin variants in the order M<Z<NHK. b) The classic ER chaperones GRP94, BiP/GRP78 and PDI contain KDEL ER localisation motifs and are detected here by anti-KDEL western blot. Their expression levels increase as predicted by the proteomic analysis, in the order M<Z<NHK expression, when resolved by subcellular fractionation using ultracentrifugation and an iodixanol gradient.

### Cellular pathways varying with terminal misfolding tendency

We next defined the sets of proteins whose expression followed the order of terminal misfolding events of α_1_-antitrypsin, i.e. positively correlated: M<Z<NHK (network analysis Fig. 2b); inversely correlated: M>Z>NHK (Fig. 2c). As the representation within these profiles was unexpectedly high relative to our initial hypothesis, we validated key findings using heatmap representations and cell biology studies (Figs 3-6).

### Translation, cytoskeleton and protein folding

Increases in terminal misfolding tendency of the expressed α_1_-antitrypsin variant (M<Z<NHK) were associated with bi-directional changes in expression of translation-associated (e.g. ribosomal proteins) and cytoskeletal proteins (Figs. 2b). Proteins associated with protein folding and re-folding were also increased in this order. The data on these were compared with heatmap colouring (Fig. 3a) and validated by Western blot data confirming altered expression of ER chaperone proteins in equally-loaded samples of cell lysate fractions (Fig. 3b). The classically-defined ER stress response pathways are not activated in CHO cell models of Z α_1_-antitrypsin expression. However, interestingly the subset of proteins associated with protein misfolding that were identified as ‘ER stress response’ do change similarly in cells expressing this and those expressing the NHK variant. Together these data indicate that the classic ER stress response may be triggered when a threshold is reached along a spectrum of misfolding-related behaviour.

### Metabolic reprogramming and mitochondrial behaviour

Proteins associated with anaerobic metabolism increased with terminal misfolding tendency (M<Z<NHK) whilst those representing aerobic respiration processes (tricarboxylic acid cycle, TCA, and oxidative phosphorylation) reduced (NHK<Z<M) as shown in Figs. 2b-c and 4a. These findings are consistent with metabolic reprogramming observed in fibrosis (47–49). Across the larger set of mitochondrial proteins in general, hierarchical cluster analysis (lower panel) confirmed that cells expressing either misfolding Z or NHK α_1_-antitrypsin resembled each other more than those expressing M (Fig. 4a). The findings were evaluated at the subcellular level in the CHO cell models. Observations were compared between the CHO cell M, Z and NHK α_1_-antitrypsin overexpression models. The functional (reactive oxidant species-producing) mitochondria in these states showed different morphological phenotypes, with elongation (reducing sphericity) increasing step-wise from CHO cell models expressing M to Z to NHK α_1_-antitrypsin (Fig. 4b). These changes were not associated with changes in Drp1 or mitofusin-1 levels (Supp. Fig. 9).

**Fig. 4:**
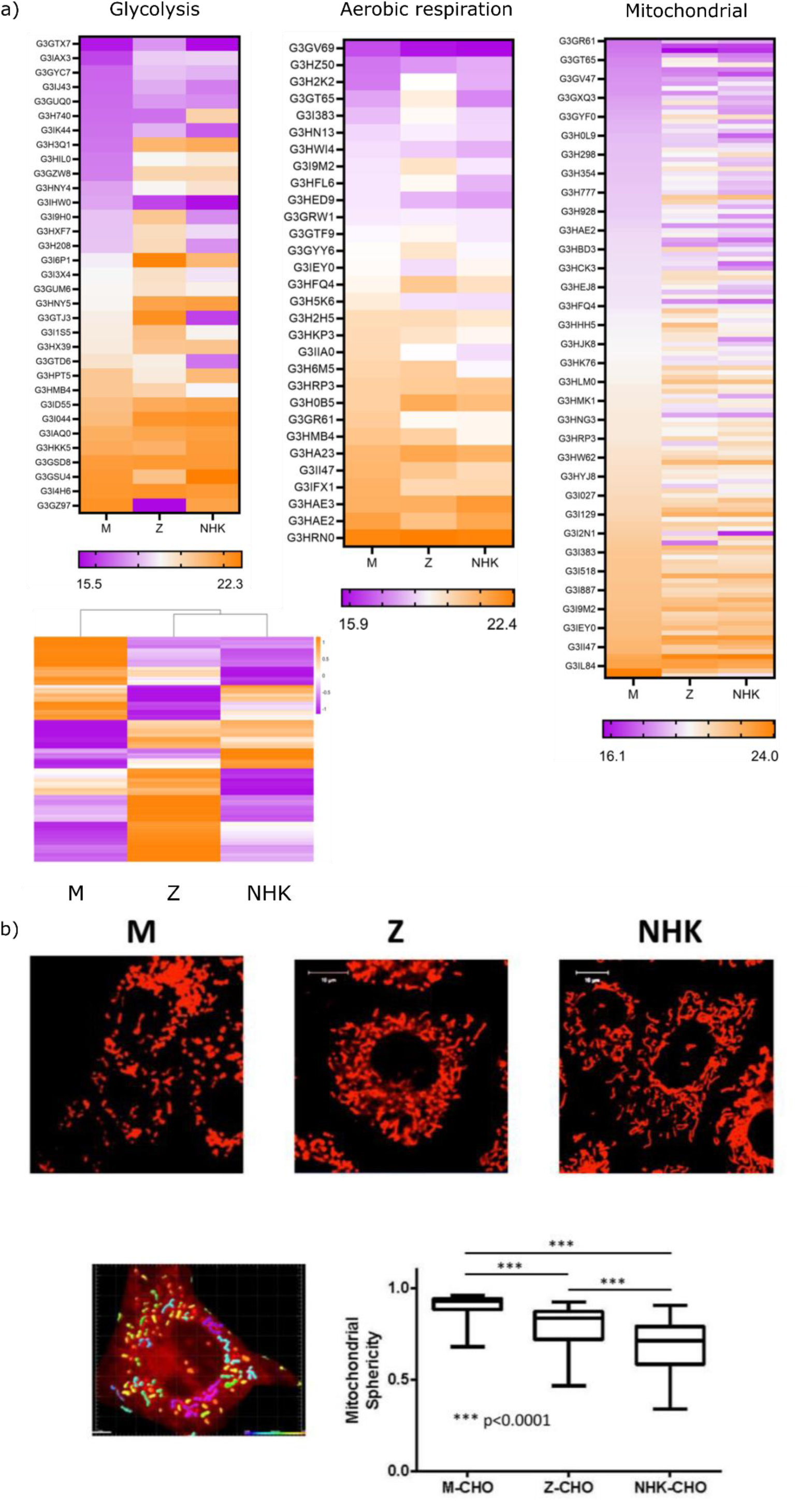
Bio-energetic and mitochondrial behaviour in CHO models. a) Heatmap depiction (format as outlined for Fig. 3) of co-expression of “Anaerobic” (e.g. glycolytic) and “Aerobic” (tricarboxylic acid, oxidative phosphorylation) metabolic pathway proteins with different α_1_-antitrypsin variants. Abundance of anaerobic pathway proteins generally increased (M<Z<NHK) whilst aerobic pathway proteins generally decreased (M>Z>NHK) with terminal-misfolding tendency. More complex patterns were seen with the global set of mitochondrial proteins (“Mitochondrial”, upper). HCA confirmed the co-expression profiles with the misfolding variants were more similar to each other than to the profile observed with M α_1_-antitrypsin expression (“Mitochondrial”, lower). b) Observed changes in morphology of functional mitochondria in association with metabolic shifts observed in Fig. 4a in representative views (confocal fluorescence microscopy), and quantified by sphericity score. Upper panel shows illustrative examples of the observed morphologies of functional mitochondria associated most closely with expression of each α_1_-antitrypsin variant. Such mitochondria are shown with red fluorescent signal in the upper panel, false coloured by sphericity score in the lower panel (red high, to blue, low – representative correlation of scoring with visible features in CHO cell expressing Z α_1_-antitrypsin) with overall sphericity data from field views quantified in box and whisker plot.

We validated these findings in inducible pluripotent stem cell (iPSC)-derived hepatocyte-like (iHep) cell models of PiMM and PiZZ behaviour derived from the same PiZZ donor with and without gene editing (50). The observed, functional mitochondria were visualised in iHeps, they again demonstrated elongation in the Z homozygous state, relative to the corrected M homozygous state (Fig. 5a). Such behaviour is seen as a response of healthy mitochondria to mitophagy, as a potential mechanism to evade incorporation into the phagolysosomal pathway (51–53). The activation of mitophagy by the deleterious effects of the Z mutation on α_1_-antitrypsin behaviour was supported by EM studies of iHep cells. These demonstrated appearances consistent with mitophagy of grossly abnormal mitochondria in PiZZ but not PiMM cells (Fig. 5b). Together our findings indicate that α_1_-antitrypsin misfolding within the ER results in mitophagy of damaged mitochondria on the one hand, and selection for functional mitochondria in more elongated species.

**Fig. 5:**
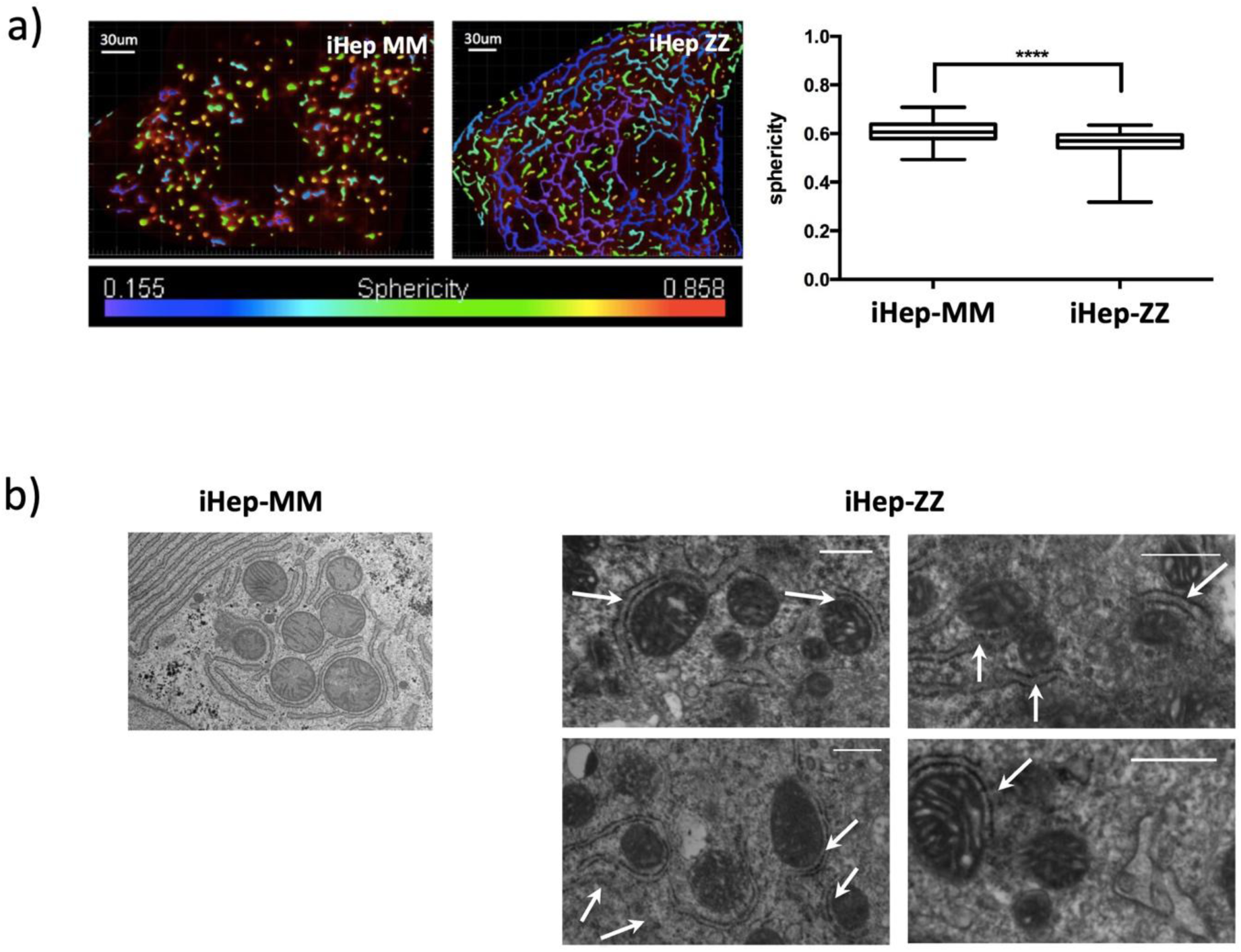
Grossly-disordered mitochondrial morphology in iHep models of Z relative to M α_1_-antitrypsin expression and ex vivo. a) Representative views of cellular mitochondria in iHep cells derived from the same donor with (MM) and without (ZZ) correction of the pathogenic α_1_-antitrypsin mutation, false coloured by sphericity score as in Fig. 4. Box and whisker plot quantitation. b) Mitochondria within MM genotype iHep model and changes seen with the PiZZ genotype cells observed by SEM, with double membrane structures (white arrows) wrapping around grossly abnormal mitochondria, consistent with mitophagy and observations ex vivo (53).

### Conserved and α_1_-antitrypsin deficiency-associated fibrotic pathways

Proteins are not annotated for specifically pro-fibrotic roles by gene ontology or network analysis descriptors. Moreover, CHO paralogs for many proteins identified in human disease pathways are not well-defined. These factors may have contributed to the low number of pro-fibrotic pathway proteins identified following a bespoke search. One notable exception was serpinH1, also known as heat shock protein (HSP)47. This protein chaperones formation of the functionally-critical triple helical structure of collagen (54), a defining component of scar tissue in all fibrosis. Such extracellular matrix proteins are generally synthesised by mesenchymal rather than cells of an epithelial lineage such as CHO cells and hepatocytes. However in fibrotic contexts epithelial cells develop mesenchymal characteristics (55). SerpinH1/HSP47 expression increased with terminal misfolding tendency of the expressed α_1_-antitrypsin variant (M<Z<NHK).

Another important set of proteins whose co-expression was associated with misfolding tendency was the integrins. α- and β-integrin expression levels increased in line with misfolding tendency (M<Z<NHK). Stepwise changes were also identified in the same direction with two important proteins that interact functionally with integrins. CD98 heavy chain (CD98hc, gene identifier SLC3A2) is constitutively complexed with integrins (56). Such complexes may stabilise the pro-fibrotic TGF-β1 signalling pathway via interactions with galectin-3 and the TGF-β receptor (TGF-βR)2 (57). Interestingly galectin-3 levels were increased in the context of NHK but not Z variant α_1_-antitrypsin expression. Levels of G-protein peptides (small monomeric GTPase protein) also increased with misfolding tendency (M<Z<NHK α_1_-antitrypsin expression models). G-proteins such as Rho, Ras and Rab proteins activate integrin functions, with Rho-mediated pathways identified as key elements of pro-fibrotic signalling (58). Levels of the alternative integrin interactors intracellular adhesion molecule (ICAM)-1 (59), and tetraspanin (60), a human paralog of which protects against fibrosis (61) showed reciprocal decreases (M>Z>NHK). Levels of talin, that links integrins with the actin cytoskeleton and hence can connect extracellular and intracellular stiffening processes (62), increased in the context of NHK though not Z α_1_-antitrypsin expression relative to expression of the wild-type M protein. To confirm pro-fibrotic behaviours suggested by the proteomic data, galectin-3:β1-integrin interactions at the cell surface were assessed in CHO cells (Fig. 6). These demonstrated increased co-localisation in the order M<Z<NHK variant expression.

**Fig. 6:**
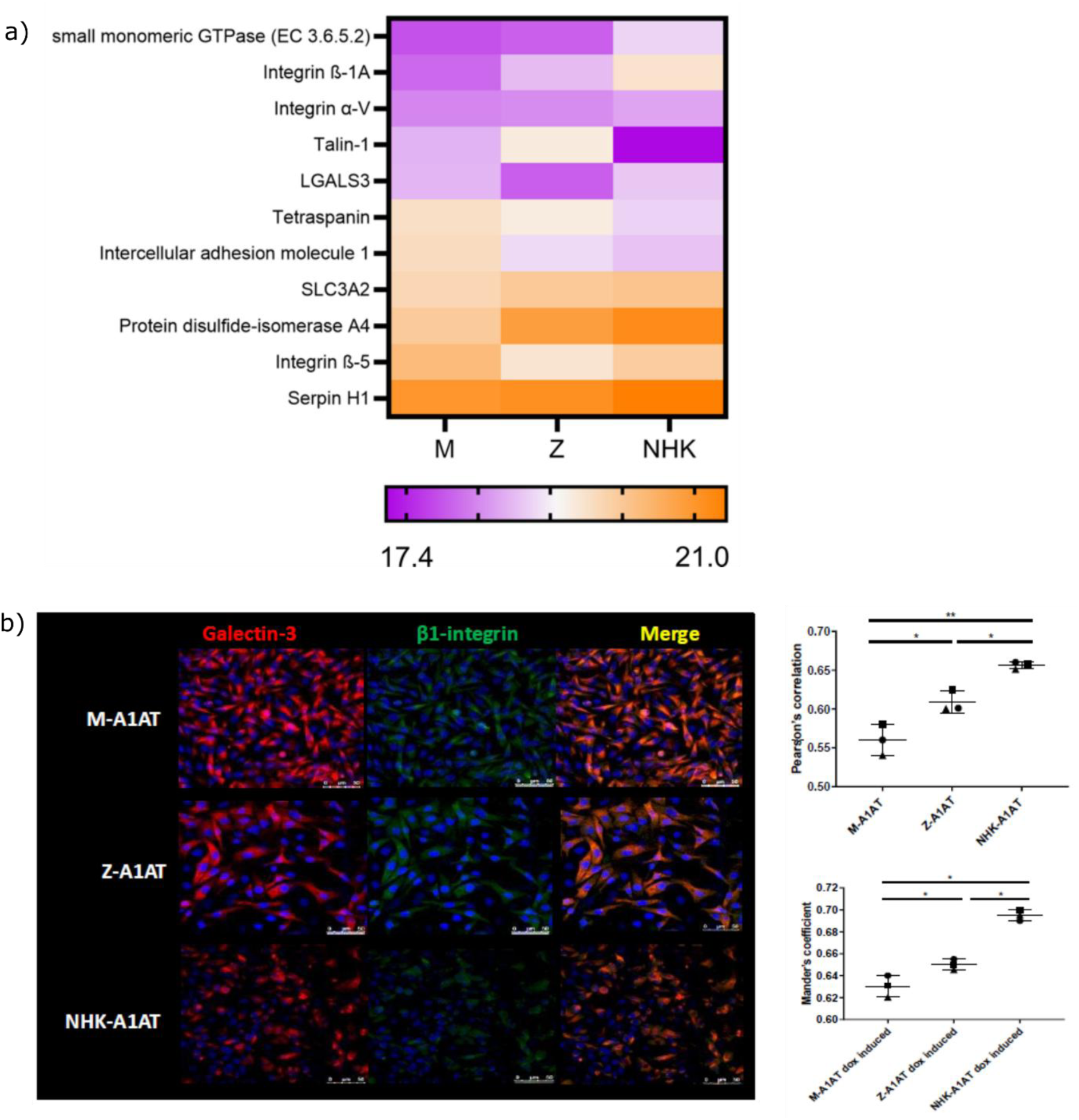
General and A1ATD-specific fibrosis mechanisms. a) Heatmap representation of co-expression levels of proteins involved in fibrosis pathways and liver disease risk in PiZZ individuals in cells expressing M, Z or NHK α_1_-antitrypsin. Formatted as in Fig. 3. b) CHO cell co-expression of galectin-3 and β1-integrin with M, Z or NHK α_1_-antitrypsin with overlapping co-immunofluorescence correlated (right panels) by Pearson’s (upper) and Mander’s (lower) co-efficients.

The increasing prevalence of protein disulphide isomerase (PDI)A4 with misfolding tendency is also of specific interest as its expression levels are correlated with liver fibrosis associated with circulating α_1_-antitrypsin deficiency in Z variant homozygotes (63). This was elevated in the context of the CHO model of Z α_1_-antitrypsin expression relative to the M expression model, but levels were higher still in the context of NHK α_1_-antitrypsin expression. Together the proteomic findings suggest that expression of Z α_1_-antitrypsin triggers intracellular responses within the ER, cytosol and cell membrane that are significantly related to terminal misfolding rather than polymerisation *per se.* The profiles of the changes between α_1_-antitrypsin variant expression models supported a role of terminal misfolding as a direct stimulator of cellular responses on-pathway for profibrotic behaviour.

### Senescence

To test these implications arising from analyses of the proteomic data, we next assessed senescence as a range of cellular behaviours associated with fibrotic disease (64). Based upon the proteomic findings, we hypothesised that multiple senescence markers would track with terminal misfolding rather than polymerisation tendencies. We therefore assessed senescence behaviours in the CHO cells expressing the different α_1_-antitrypsin variants (Fig. 7). Again, these demonstrated stepwise changes supporting the hypothesis. Senescence-associated-β-galactosidase (SA-β-gal) staining increased in cells expressing Z α_1_-antitrypsin, relative to those expressing M α_1_-antitrypsin, and increased still further in cells expressing the NHK variant (Fig. 7a). As noted, mitochondrial dysfunction, that is regarded as a marker of senescence, also appears to increase in the same order Consistent with this, proliferation assessed by BrdU assay reduced stepwise with expression of M to Z to NHK α_1_-antitrypsin (Fig. 7b). Interestingly, however, p16 staining was unchanged between cells expressing the three variants (Fig. 7c). Together with the evidence of mitochondrial morphologic changes that correlated with dysfunctional species (Fig. 5b), also regarded as a marker of senescence behaviour, these findings indicated that senescence was occurring to extents that varied with the degree of terminal misfolding in these cell models. However there was no increase in late phase senescence behaviour as reported by p16 expression within the timeframe of the experiment.

**Fig. 7:**
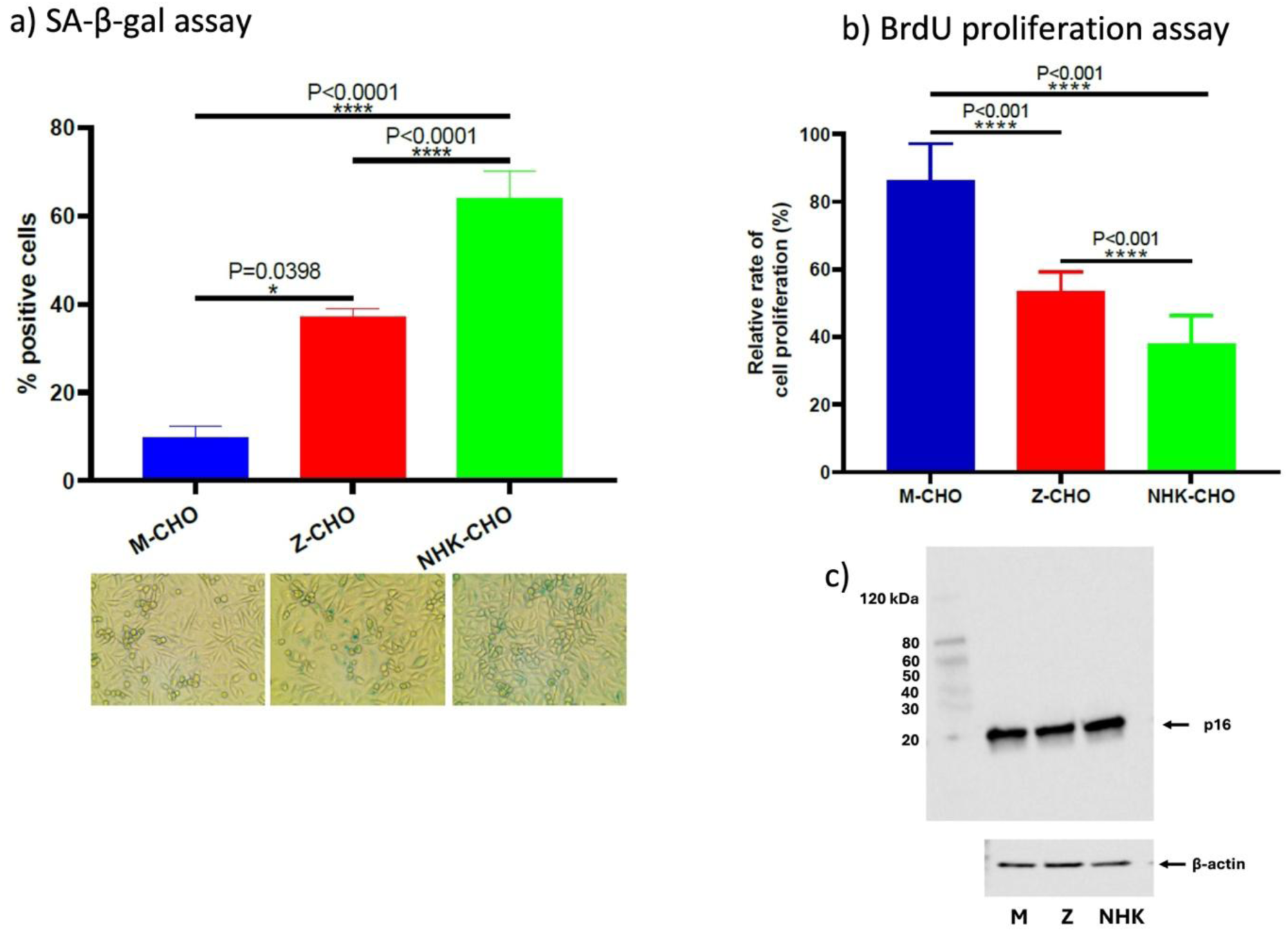
Senescence readouts. assessed for CHO cells expressing M, Z or NHK α_1_-antitrypsin, n=3. a) SA-β-galactosidase stain intensity quantitation (above) and representative micrographs (lower panel). Increases indicate increased senescence behaviour. b) BrdU proliferation assay, reductions indicate increased senescence behaviour. c) Western blot analysis of SDS-PAGE, probing for p16 (upper panel, molecular mass markers in left hand lane as indicated) and β-actin loading control (lower pane, equal volume samples from same lysate run on separate gel).

## Discussion

Alpha_1_-antitrypsin deficiency-associated liver disease results from abnormal structural behaviour of the α_1_-antitrypsin polypeptide, rendering it vulnerable to terminal misfolding and ERAD, and in parallel to the formation of α_1_-antitrypsin polymers. The hepatic fibrosis has been directly attributed to the observed accumulation of the latter within hepatocyte inclusion bodies. However the findings in this paper indicate that terminal misfolding of α_1_-antitrypsin can itself drive phenotypic changes in cell models of α_1_-antitrypsin that are on-pathway for fibrosis. Specifically, we have identified metabolic reprogramming, senescence and cell surface markers of pro-fibrotic responses. This is surprising as expression of the Z variant in CHO cells, iHeps and hepatocyte models is not associated with the unfolded protein response (UPR) or ER stress observed with expression of the NHK variant in cell models (20).

We and others have previously reported changes in metabolic pathways in iHeps expressing Z rather than M α_1_-antitrypsin (33, 65). Metabolic reprogramming is a hallmark of fibrotic behaviour in pathological fibrosis in the lung and other organs (47–49). It is characterised by a shift from oxidative metabolic pathways to glycolytic behaviour. However, our studies are the first to suggest that in α_1_-antitrypsin deficiency these may correlate with the terminally-misfolded protein load rather than solely from the polymer burden.

In our studies, this was associated with a change in mitochondrial morphology. Changes in mitochondrial sphericity are described in other protein misfolding disorders and even in models of dementia caused by misfolding and polymerisation of neuroserpin, a protein closely related to α_1_-antitrypsin (66). Commonly, the pathological state is associated with fragmentation of mitochondria to smaller and hence more spherical states as a precursor to apoptosis. Studies of other Z homozygote iHep models of α_1_-antitrypsin deficiency report similar findings to this (65). Our EM cellular studies presented here and previously (33) also demonstrated this behaviour. Conversely the findings from our fluorescence studies report increased elongation of observed species across large numbers of mitochondria. These differences are explained by the choice of fluorescent dye used in the studies, and hence what is reported. The studies by Kaserman *et al* used a dye that stains the mitochondrial membrane and will therefore identify all mitochondrial species (65). For our fluorescence work following up the proteomic data on bioenergetic pathways we were particularly interested in the appearances of functional mitochondria. We therefore used a marker for this, i.e. a dye that recognises non-depolarised species (preserved Δψm). The morphologies of these mitochondria visualised in M and Z CHO cells were validated by the observation of similar phenotypes in the corresponding iHep models. However the EM appearances in the latter indicate severe co-existent mitochondrial degeneration with closely-associated evidence of early mitophagy in PiZZ compared with PiMM iHeps. These characteristics define depolarised, dysfunctional mitochondria that will not be reported with our fluorescent marker. Taken together, these findings very closely recapitulate observations of mitochondrial injury, mitophagy and elongation observed in PiZZ liver tissue with established liver disease (53). Mitochondrial dysfunction (67) and megamitochondria (68) are also observed as common features in a range of chronic liver diseases. The findings in this study demonstrate that in α_1_-antitrypsin deficiency such changes may be early events, closely coupled with protein misfolding, but not specifically to polymerisation, within the ER.

We note that translocation of aggregated Z α_1_-antitrypsin into mitochondria has been observed in a liver-derived cell line (69). Our studies were not designed to look for this. If this were indeed polymeric material, it should not be centrally relevant to the process that we are observing not only with Z α_1_-antitrypsin but also to a greater degree with expression of the NHK variant which cannot polymerise.

To explore further the possibility that terminal misfolding of α_1_-antitrypsin rather than polymerisation alone correlates with cellular processes that are on-pathway for liver fibrosis we went on to evaluate markers of senescence in these cell lines. The findings indicated increased senescence behaviour with increasing misfolding, again following the pattern M<Z<NHK variant expression. Lastly, we looked at the cell surface co-localisation of galectin-3 and β1-integrin. Both proteins are common mediators of fibrosis across multiple organs including lung and liver (57, 70, 71). We have identified an increase in their close (<40 nm) co-localisation at the cell surface in epithelial cell and tissue models of early lung fibrosis in response to stimulation by the key pro-fibrotic cytokine transforming growth factor (TGF-)β1 (57). It is therefore interesting to observe similar close co-localisation in response to expression of the NHK variant compared to M α_1_-antitrypsin in the CHO cell models of liver fibrosis, and that this is also observed to a lesser extent with expression of the Z mutant.

We have therefore identified a range of responses to expression of pathogenic α_1_-antitrypsin mutants that can polymerise and/or undergo terminal misfolding, that increase as the latter increases, even when polymerisation cannot be achieved. From the proteomic to subcellular and functional levels these are generally on-pathway for fibrosis. Fig. 8 outlines two ways in which these could arise within the different models. Firstly, it is possible that the common patterns of cellular changes reported here at multiple scales with both misfolding variants could have different triggers in each case. With the Z α_1_-antitrypsin expression models, this may be the formation of polymers but not the UPR and ER stress response, since these models do not manifest this in the absence of a further proteostatic challenge (“second hit”). Conversely, in the NHK α_1_-antitrypsin expression model where polymerisation cannot occur, but the UPR and ER stress response are activated, the latter would be the likeliest triggers.

**Fig. 8:**
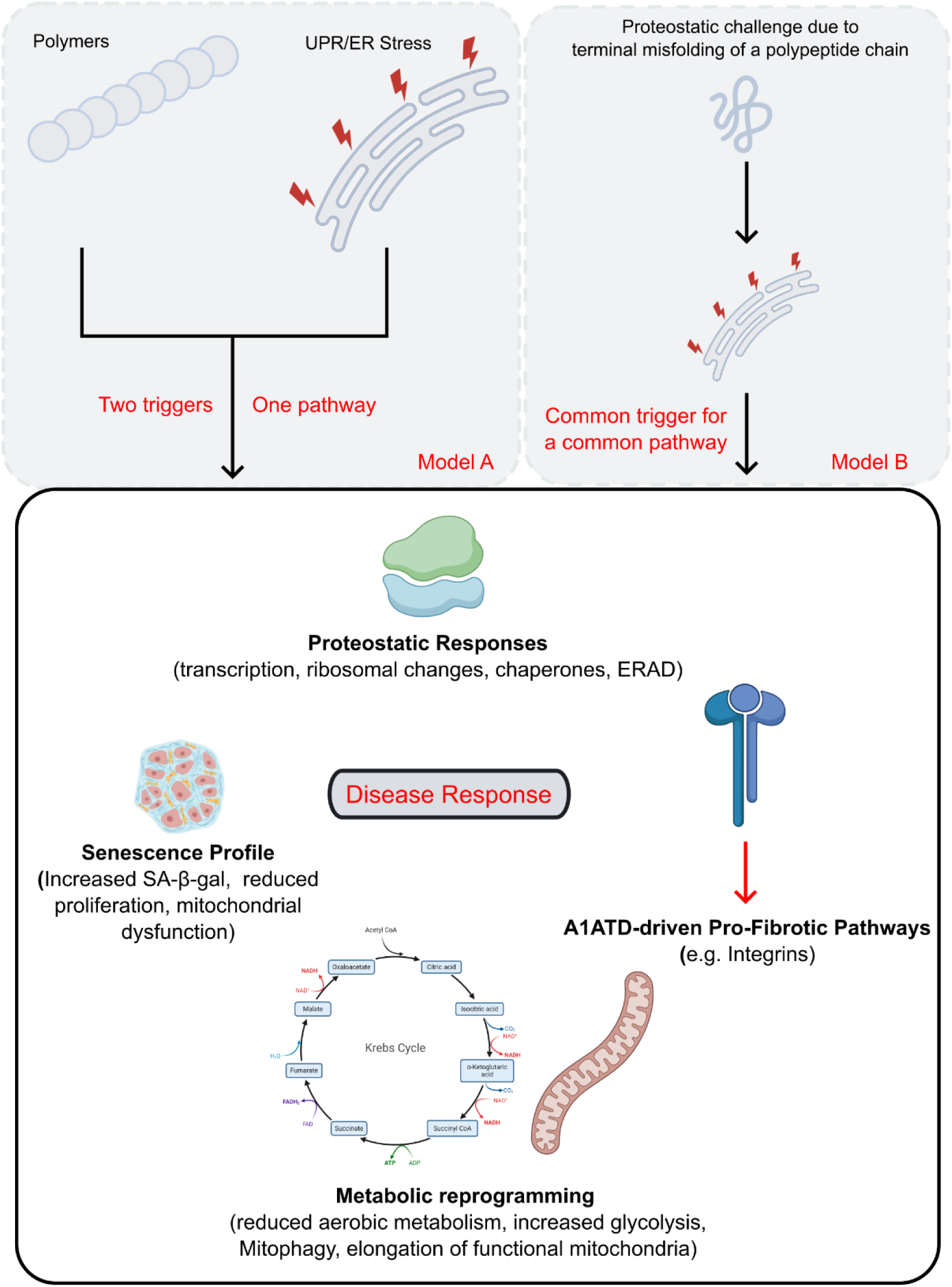
Different models for the activation of pro-fibrotic behaviour. Model A includes the combination of polymer accumulation and UPR/ER stress to drive a two-trigger pathway. Model B involves the stimulation of pro-fibrotic activity from a common primary trigger, the terminally-misfolded fraction of α_1_-antitrypsin that is recognised and targeted for degradation by ERAD.

The second hypothesis is that the common responses described here arise from a common primary trigger, the terminally-misfolded fraction of α_1_-antitrypsin that is recognised and targeted for degradation by ERAD. This has been quantified as the fate of ∼70% of all Z α_1_-antitrypsin polypeptides synthesised and will account for 100% of synthesised NHK variant polypeptides (4). This is the more parsimonious explanation. It complicates previous understanding of polymerisation as the key toxic gain-of-function event in α_1_-antitrypsin deficiency-associated liver disease (24), and the significance of some identified polymer-response pathways (72). On the other hand it is highly consistent with the priming of PiZZ disease models to induction of the UPR and ER stress in response to a second hit, relative to the PiMM state (19, 73). This hypothesis is also consistent with findings from hypothesis-free (exomic and proteomic) studies of the effects of the Z mutation in cell models and human subjects (63, 74). These most strongly implicated ER proteins involved in the general misfolding (protein disulphide isomerase (PDI)A4) and ERAD responses as mediators of liver toxicity and fibrosis in PiZZ α_1_-antitrypsin deficiency.

The two mechanistic hypotheses need not be entirely mutually exclusive, and the contribution of terminal misfolding relative to polymerisation loads in liver disease associated with PiZZ α_1_-antitrypsin deficiency deserves further clarification. The findings further indicate that the homozygote states for the NHK and other variants of α_1_-antitrypsin where terminal misfolding causes a Null phenotype should not be assumed benign in terms of liver disease risk. They may therefore merit increased clinical vigilance, and advice on minimising ‘second hits’ such as excessive alcohol intake or metabolic syndromes. The absence of an observed clinical association to date may relate to their rarity, rather than representing evidence of absence of cellular pathology.

## Acknowledgements

PS received support from the National Institute for Health Research (NIHR) Leicester Biomedical Research Centre (BRC) Respiratory Theme, British Lung Foundation (BLF)/Asthma+Lung (A+L)UK and the Wellcome Trust (WT). KRA and KT have received support from the WT and Alpha-1 Foundation (A1F). MPN was supported by A1F and a WT VIP award. AC was supported by Waters Ltd and a BBSRC CASE studentship. NOL was supported by an MRC studentship. IH, JAI, DAL were supported by the Medical Research Council (UK, MRC), WT, GlaxoSmithKline, the Rosetrees Trust, and the NIHR UCLH BRC. C-PS was supported by the Children’s Liver Disease Foundation. GS was supported by a Wellcome Trust ISMB DTP studentship. AO was supported by an eALTA award. JB was supported by an MRC AIM PhD studentship. AWEJ was supported by the MRC, the Australian National Health and Medical Research Council, the Australian Mitochondrial Disease Foundation and Victorian Government Operational Infrastructure Support. STR was supported by WT, MRC, NIHR Imperial BRC, A1F, and NC3Rs. LV was supported by European Research Council (ERC), NIHR CUH BRC, and the WT/MRC Cambridge Stem Cell Institute. BG received support from NIHR Leicester BRC Respiratory Theme, BLF/A+LUK, MRC and A1F.

We are grateful to Prof Stefan Marciniak (University of Cambridge) for very helpful discussions in the development of this work. We thank the Advanced Imaging Facility (RRID:SCR_020967) at the University of Leicester for support.

---

**Supp. Fig. 1**

**Cell free biophysics of polymerisation and misfolding, including Alpha-1 Antitrypsin variants M, Z, S, Baghdad and NHK.**

**Supp. Fig. 2**

**Qualitatively similar changes observed in CHO cells expressing NHK and Z relative to findings with M α_1_-antitrypsin expression but not following order of terminal misfolding or polymerisation tendencies.**

Increasing expression with M<NHK<Z (a) and Z<NHK<M (b).

**Supp. Fig. 3-8**

Pathway analyses for all profiles shown in Fig. 2 and Supp. Fig. 2.

**Supp. Fig. 9**

SDS-PAGE of cell lysates for CHO cell models expressing M, Z and NHK α_1_-antitrypsin probed with western blots using antibodies to recognize Drp1 and Mfn-1. Sample loading demonstrated by β-actin loading shown below.

## Notes

### Competing Interest Statement

The authors have declared no competing interest.

### Summary of Updates

Figures now available after technical issue solved.

