## Supplementary material for "Both terminal misfolding and polymerisation contribute to disease-relevant responses in cell models of α_1_-antitrypsin deficiency-associated liver disease": Figures and Supplementary

[6] Universidad Católica de Murcia (UCAM), HiTech. Campus de los Jerónimos 135, E-30107, Guadalupe, Murcia, Spain.

[7] Department of Women and Children's Health, School of Life Course Sciences, King's College London, London, UK;

[8] Department of Metabolism, Digestion and Reproduction, Faculty of Medicine, Imperial College, Commonwealth Building, Du Cane Road, London, W12 0NN, UK;

[9] Berlin Centre for Regenerative Therapies (BCRT), Berlin Institute of Health (BIH), Charité – Universitätsmedizin, and Max Planck Institute for Molecular Genetics, Berlin, Germany.

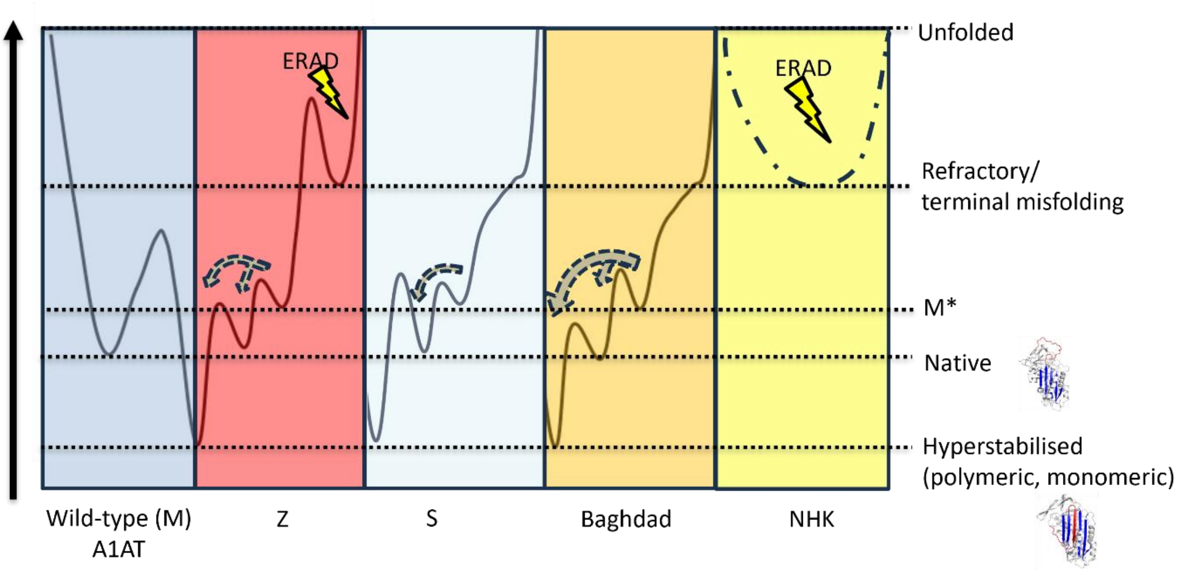


**Supp. Fig. 1**: Cell free biophysics of polymerisation and misfolding, including Alpha-1 Antitrypsin variants M, Z, S, Baghdad and NHK.

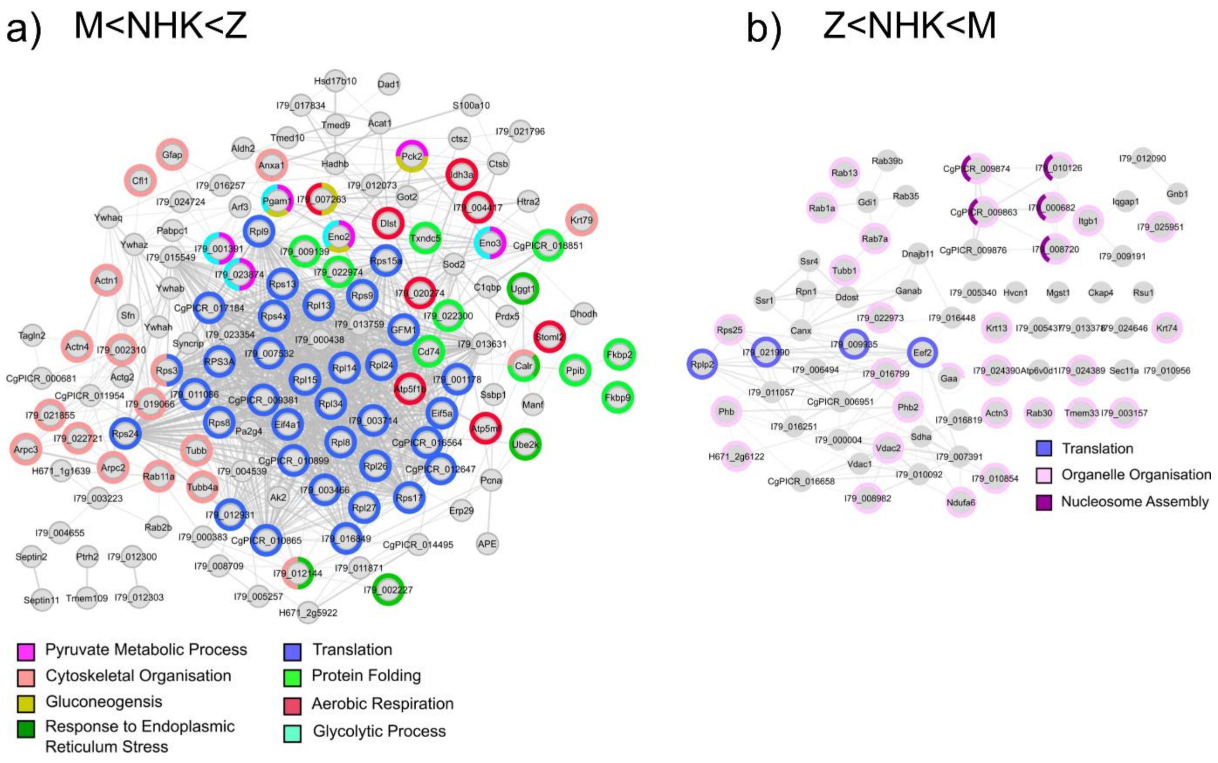


**Supp. Fig. 2**: Qualitatively similar changes observed in CHO cells expressing NHK and Z relative to findings with M α$_1$-antitrypsin expression but not following order of terminal misfolding or polymerisation tendencies.

Increasing expression with M<NHK<Z (a) and Z<NHK<M (b).

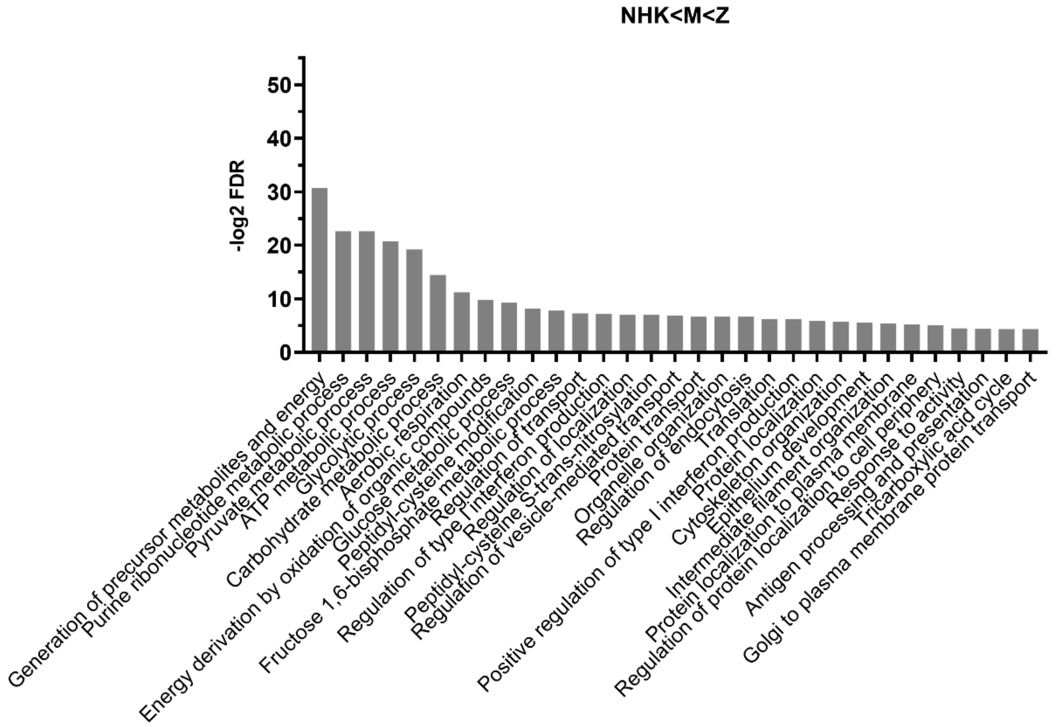


**Supp. Fig. 3**: Gene Ontology Analysis including biological processes and molecular functions, for proteins that followed the abundance trend NHK<M<Z.

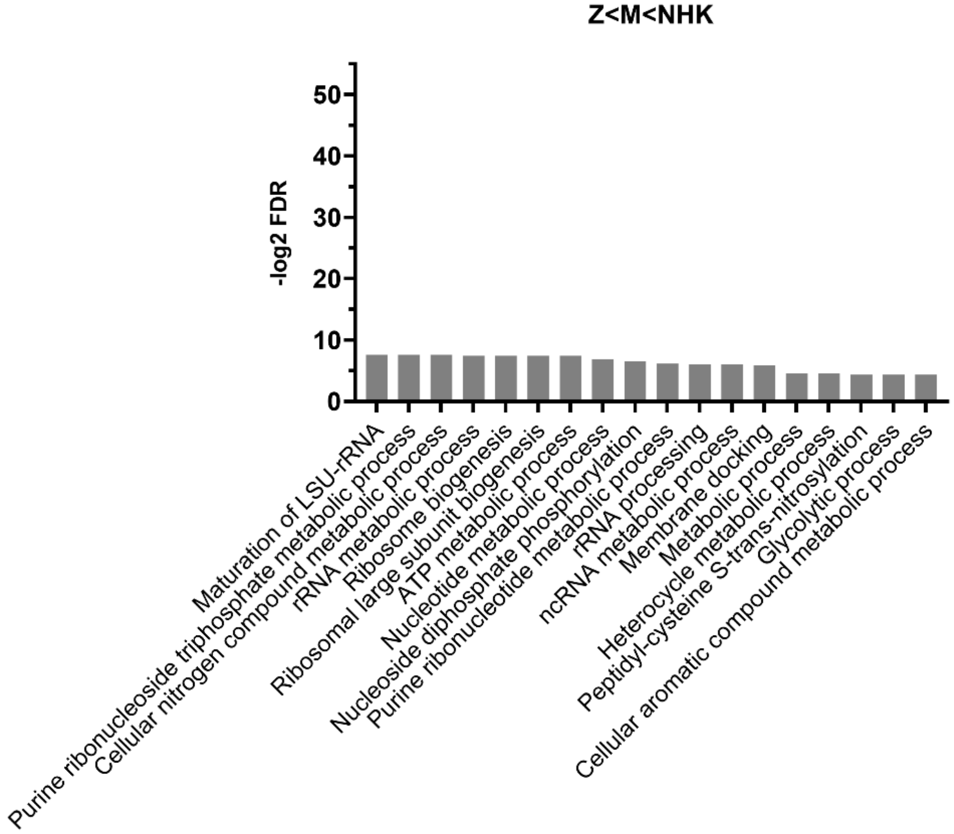


**Supp. Fig. 4**: Gene Ontology Analysis including biological processes and molecular functions, for proteins that followed the abundance trend Z<M<NHK.

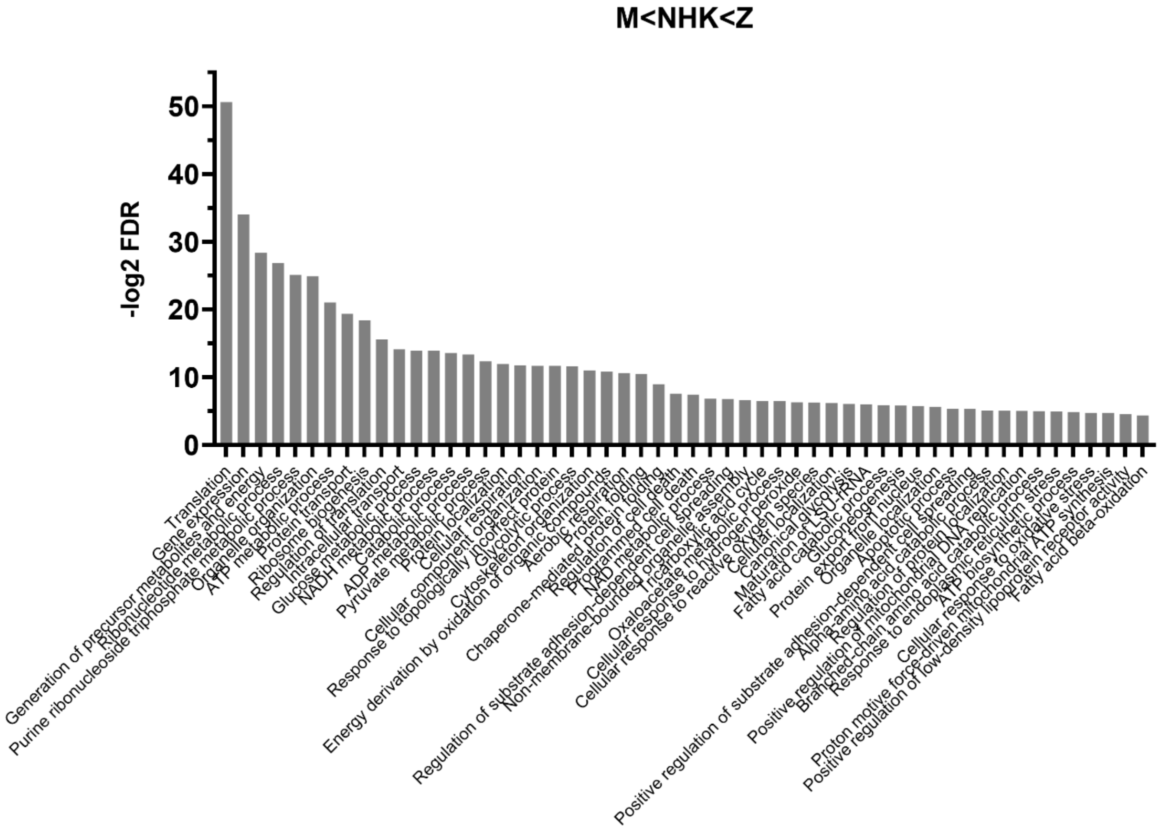


**Supp. Fig. 5**: Gene Ontology Analysis including biological processes and molecular functions, for proteins that followed the abundance trend M<NHK<Z.

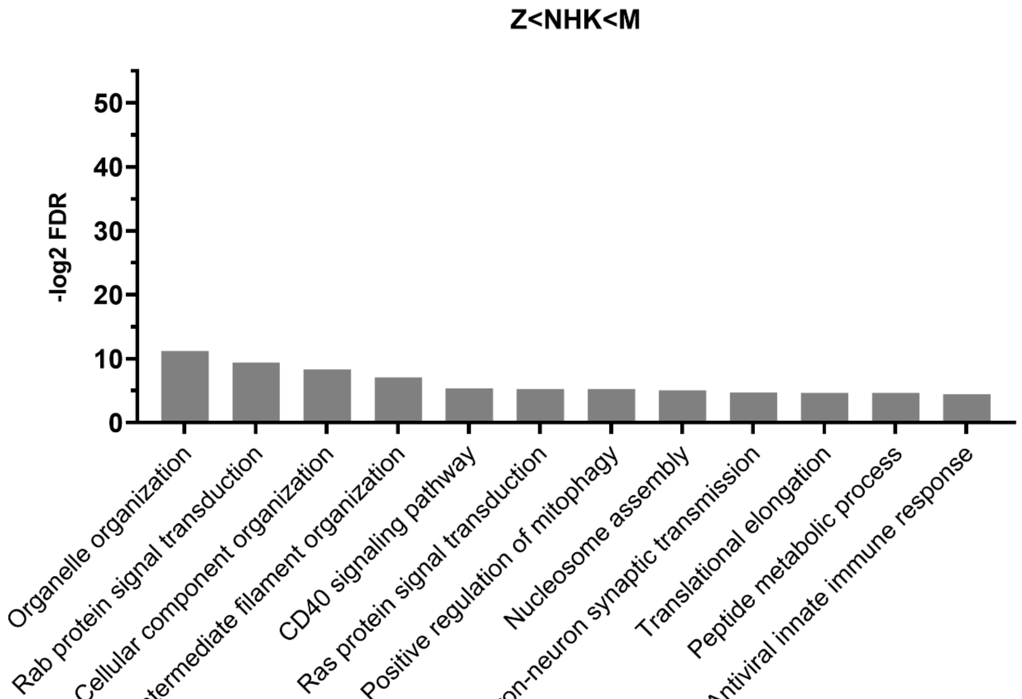


**Supp. Fig. 6**: Gene Ontology Analysis including biological processes and molecular functions, for proteins that followed the abundance trend Z<NHK<M.

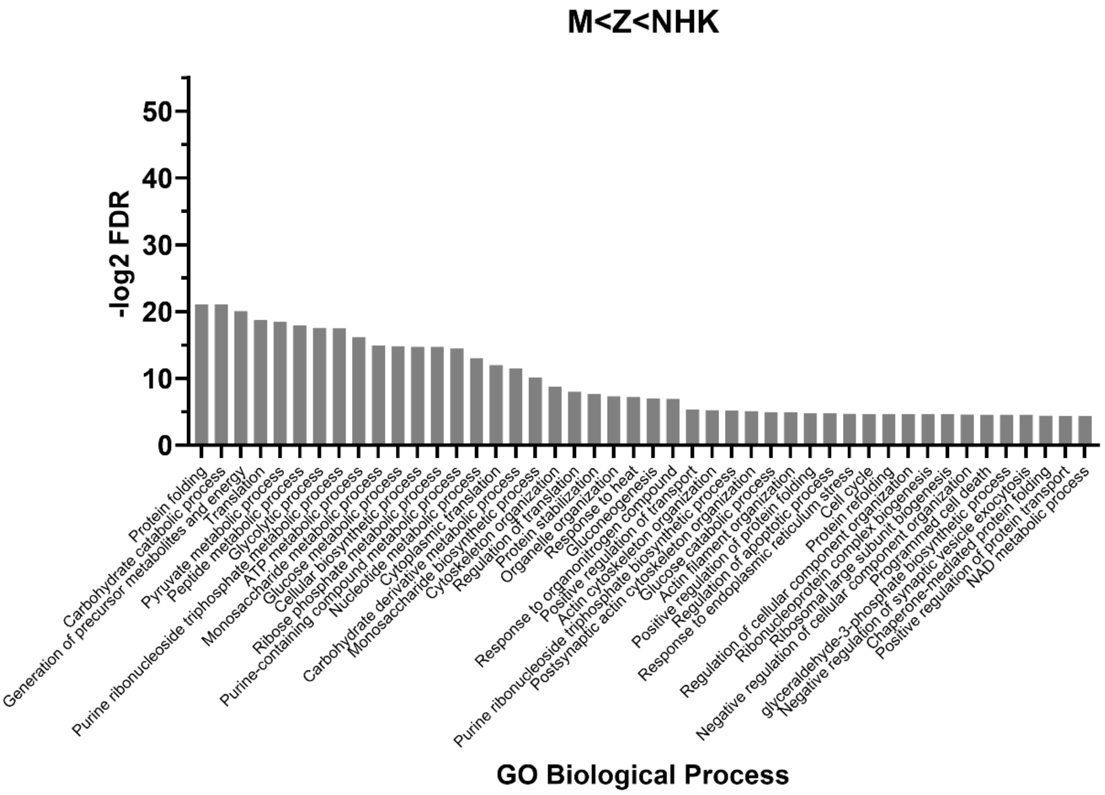


**Supp. Fig. 7**: Gene Ontology Analysis including biological processes and molecular functions, for proteins that followed the abundance trend M<Z<NHK.

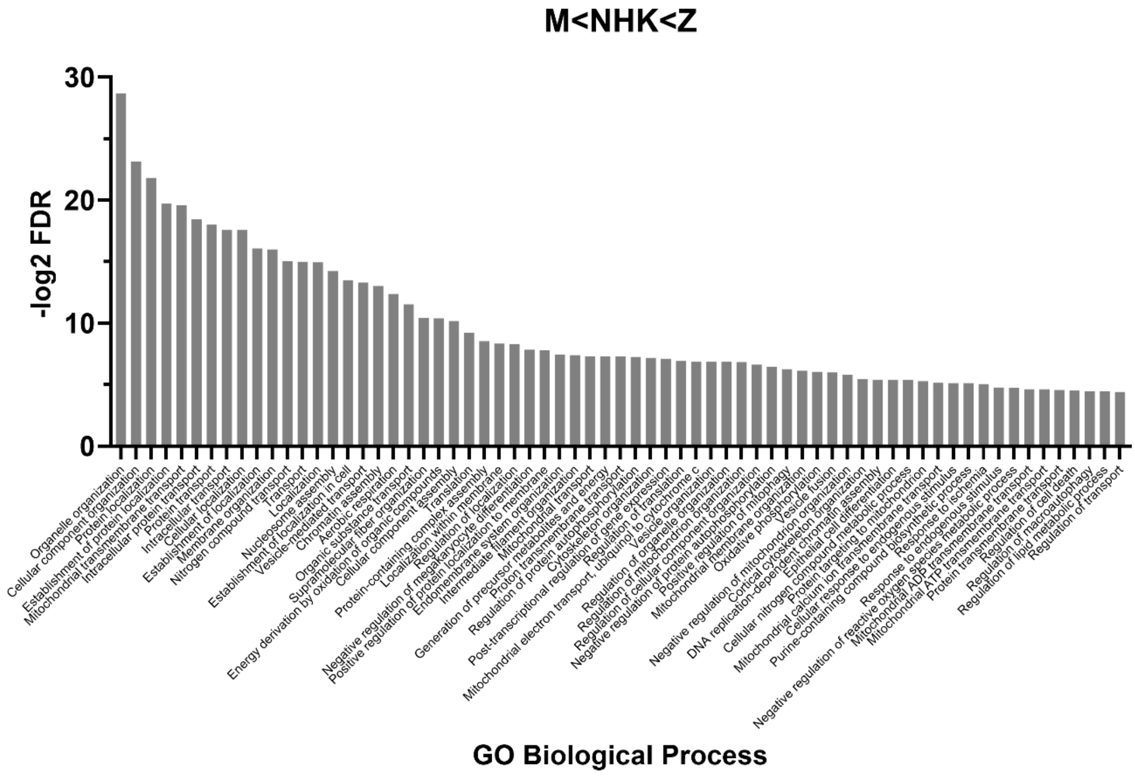


**Supp. Fig. 8**: Gene Ontology Analysis including biological processes and molecular functions, for proteins that followed the abundance trend M<NHK<Z.

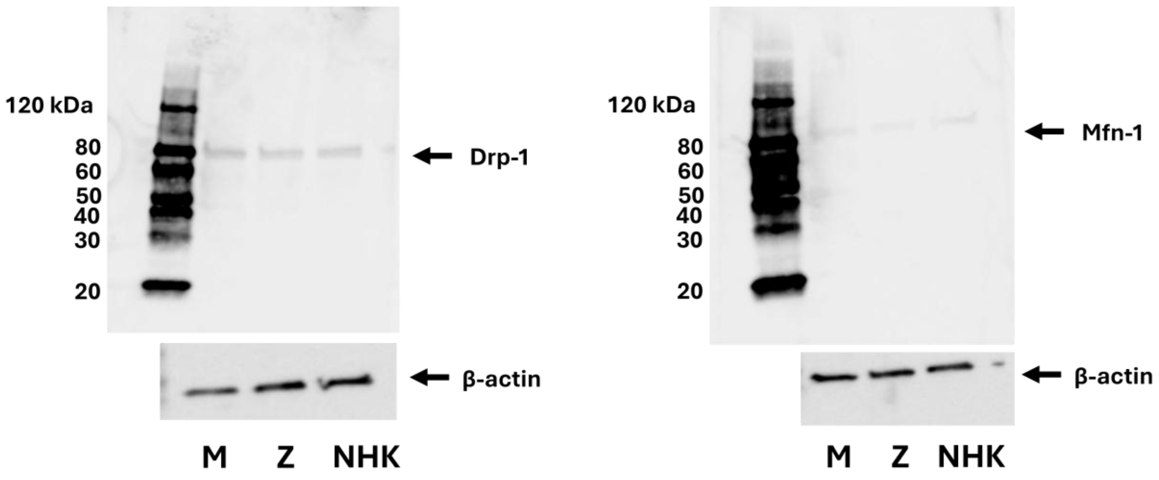


**Supp. Fig. 9**: SDS-PAGE of cell lysates for CHO cell models expressing M, Z and NHK $\alpha_1$-antitrypsin probed with western blots using antibodies to recognize Drp1 and Mfn-1. Sample loading demonstrated by β-actin loading shown below.
